# Characterizing physical interactions between male and female mosquitoes (Aedes aegypti) in relation to female receptivity and insemination outcomes using a hydrophobic fluorescent dye

**DOI:** 10.1101/2023.05.25.542180

**Authors:** Monica M. Cramer, Thomas M. Gabel, Laura B. Duvall

## Abstract

*Aedes aegypti*, the yellow fever mosquito, presents a major threat to human health across the globe as a vector of disease-causing pathogens. Females of this species generally mate only once. From this single mating event, the female stores sufficient sperm to fertilize the multiple clutches of eggs produced during her lifetime. Mating causes dramatic changes in the female’s behavior and physiology, including a lifetime suppression of her mating receptivity. Female rejection behaviors include male avoidance, abdominal twisting, wing-flicking, kicking, and not opening vaginal plates or extruding the ovipositor. Many of these events occur on a scale that is too miniscule or fast to see by eye, so high-resolution videography has been used to observe these behaviors instead. However, videography can be labor intensive, require specialized equipment, and often requires restrained animals. We used an efficient, low-cost method to record physical contact between males and females during attempted and successful mating, determined by recording spermathecal filling after dissection. A hydrophobic oil-based fluorescent dye can be applied to the abdominal tip of one animal and can be subsequently transferred to the genitalia of animals of the opposite sex when genital contact occurs. Our data indicate that male mosquitoes make high levels of contact with both receptive and unreceptive females and that males attempt to mate with more females than they successfully inseminate. Female mosquitoes with disrupted remating suppression mate with and produce offspring from multiple males, transferring dye to each. These data suggest that physical copulatory interactions occur independently of the female’s receptivity to mate and that many of these interactions represent unsuccessful mating attempts that do not result in insemination.

## INTRODUCTION

*Aedes aegypti*, the yellow fever mosquito, is a global public health concern as a vector of disease-causing pathogens, including yellow fever, dengue fever, chikungunya, and Zika (Rogers et al. 2006; Bhatt et al. 2013; Guerbois et al. 2016; Weaver et al. 2016). Females of this species are obligate blood feeders; to successfully reproduce, they need to mate and consume blood to obtain necessary protein for egg development (Dimond et al. 1956; Attardo et al. 2005). Although limited remating has been reported across *Ae. aegypti* populations, this species is generally monandrous – females mate only once (Craig 1967; Gwadz et al. 1971; Jones and Pilitt 1973). From this single mating event, a female stores sufficient sperm to fertilize all of the eggs produced across multiple clutches for the rest of her life (Craig 1967; Spielman et al. 1967; Carvalho et al. 2018). Specialized organs, called spermathecae, store and maintain sperm after transfer to the female, which can then be released when eggs are ready to be fertilized (Roth 1948). Mating causes dramatic changes in female behavior, including the lifetime suppression of receptivity, inducing her to reject all subsequent mating attempts (Hiss and Fuchs 1972; Clifton et al. 2014). Methods to assess male-female interactions, including attempted mating, successful mating, and rejection, are of great interest to researchers studying *Ae. aegypti* behavior. A better understanding of the mechanisms by which *Ae. aegypti* mate will facilitate their exploitation for new population control strategies by preventing mating and reproduction.

Early examinations of *Ae. aegypti* mating behavior characterized females as highly promiscuous, based on the frequency with which mated females appear to copulate with subsequent males (MacGregor 1915; Roth 1948). However, later studies using genetic markers of offspring paternity showed that *Ae. aegypti* females become refractory to further mating after successful insemination (Craig 1967; Spielman et al. 1967), and reject future prospective mates by male avoidance, kicking, preventing a male from assuming the correct position, abdominal twisting to prevent copulation, and/or failing to extrude her ovipositor (Roth 1948; Gwadz et al. 1971; Jones and Pilitt 1973; Cator and Harrington 2011). Thus, some forms of female rejection may be indiscernible from successful mating to the naked eye (Eberhard 1991).

This phenomenon, by which male-derived mating signals prevent a female from further mating, is referred to as paternity enforcement. Short-term paternity enforcement within hours of mating is regulated in part by a peptide found in male seminal fluid, Head Peptide (HP-I), which activates a cognate receptor in females, Neuropeptide Y-Like Receptor 1 (NPYLR1) (Naccarati et al. 2012; Duvall et al. 2017). Females mutant for *npylr1* are receptive to subsequent mates, bearing mixed-paternity offspring when exposed to successive males within hours. *Hp-I* mutant males fail to enforce short-term paternity, fathering only a portion of his mate’s offspring if she remates with a subsequently-encountered male within hours, although slower acting paternity enforcing signals are still present in *hp-I* mutant males (Duvall et al. 2017). The signals that induce lifetime paternity enforcement and the mechanisms that maintain this behavioral change in females remain unknown (Craig 1967; Fuchs et al. 1968; Hiss and Fuchs 1972).

Assessing the mating status of *Ae. aegypti* females typically requires dissecting the female reproductive tract to assess the presence of sperm in two of the three spermathecae, although it is also possible to score insemination in a live, immobilized female (Carrasquilla and Lounibos 2015). However, scoring spermathecae for insemination does not capture attempts made, or the identities of animals involved in attempted or successful mating. Because many rejection behaviors occur on a scale too small and fast to see by eye, high-speed, high-resolution videography has been used to observe males mating with tethered females whose scutellum has been glued to a pin to restrict their movement (Aldersley and Cator 2019). While videography can offer high-resolution, detailed observation of mating behavior, it can also be costly, low throughput, and requires restrained animals.

In this study, we utilized an efficient, low-cost method to assay physical contact between males and females during attempted and successful mating. A hydrophobic oil-based fluorescent dye applied to the abdominal tip of one animal is subsequently transferred to the genitalia of animals of the opposite sex during copulation. Females can be scored for insemination to distinguish between attempted and successful mating. Because mating attempts are frequent and do not always result in successful insemination, and failed mating attempts can involve contact between the animals’ abdominal tips that is indistinguishable to the naked eye from successful mating, dye transfer paired with spermathecal dissections can identify females involved in successful and unsuccessful mating attempts as well as males who attempted to mate and were not rejected before making abdominal contact. Our findings reveal males’ propensity to initiate mating regardless of female receptivity, and suggest that males come sufficiently close to mating to make genital contact but are unable to inseminate the female.

Dye applied to either males or females is transferred to the abdominal tip of animals of the opposite sex during successful mating that results in insemination, and also during unsuccessful mating attempts in which the abdominal tips come into contact. Since successful insemination requires contact between the mating pair, we do not expect to see any instance of insemination without dye transfer and did not observe any such occurrences. Previous work suggests that many mating attempts are unsuccessful (Aldersley and Cator 2019) and that males attempt to mate with unreceptive females. Because dye transfer reports both successful and unsuccessful attempts in which genital contact is achieved, we expected rates of dye transfer to exceed rates of insemination.

Our experimental methods for dye application and scoring are provided in the “Methods” section. In the “Results” section, we present our experiments showing that dye applied to either males or females is transferred to the abdominal tip of animals of the opposite sex during successful mating that results in insemination, and also during unsuccessful mating attempts in which the abdominal tips come into contact. In the “Discussion” section, we discuss our results and avenues for future research.

## METHODS

### Rearing

*Aedes aegypti* wild-type laboratory strain (Orlando) and mutant strain (*npylr1*-/-, (Liesch et al. 2013)) were reared in environmental rooms at 70 - 80% relative humidity, 25 – 28°C with a photoperiod of 14 hours light: 10 hours dark, as described in DeGennaro et al. 2013. Eggs were hatched in hatch broth (1 crushed TetraMin fish food tablet in 1L deionized, deoxygenated water). Larvae were fed crushed TetraMin fish food tablets. The *npylr1* mutant strain was selected for experiments in which females encounter successive males because these mosquitoes lack functional NPYLR1 and the females are receptive to remating but males of this strain do not show any mating deficits (Duvall et al. 2017).

For assays involving unmated females, animals were separated by sex as pupae to ensure the unmated status of the females. Unmated males and females were housed separately until behavioral assays were performed. For assays involving mated females, males and females were allowed to cohabit as adults for at least 5 days to ensure that females were mated. Adult mosquitoes were housed in custom cages (216 mm diameter, 181 mm height) and provided access to 10% sucrose *ad libitum*. For all assays, males and unmated females were 5 - 14 days post-eclosion and mated females were 7 - 21 days post-eclosion at the beginning of the assay.

### Dye Application and Scoring

Mosquitoes were cold anesthetized at 4°C for 10 minutes prior to dye application, then placed in plastic cups on ice. Using forceps, mosquitoes were transferred from the plastic cups to a chilled glass petri dish, where fluorescent oil-based dye (ACDelco 1148963 GM Original Equipment 10-5045 Multi-Purpose Fluorescent Leak Detection Dye) was applied to the terminal two segments of the abdomen. Dye application was performed under a Leica MZ10 F Fluorescence Stereomicroscope in the Cy3 channel (Figure 1A) using a modified fine-tip paintbrush (Amazon Catalog# B073YDKWWP). Prior to dye application, all but 20 bristles were removed from a fine-tip paintbrush to minimize the amount of dye applied. After dye application, animals were placed on ice in a plastic cup lined with a Kim-wipe to absorb any excess dye and allowed to recover for approximately 5 minutes. To confirm that dye transfer reported direct physical interactions between animals, we performed control assays in which 10 painted females were housed in a custom cage (216 mm diameter, 181 mm height) for 10 hours, then removed. A new set of 10 unpainted female mosquitoes was then housed in the same cage for 12 hours, then scored for the presence of dye to confirm that no dye was transferred incidentally via cage surfaces (Supplemental Data Table 1). To confirm that dye was not transferred between female mosquitoes during non-mating interactions, dye was applied to 10 unmated female mosquitoes which were cohoused with 10 unmated, unpainted female mosquitoes for 20 hours, after which all animals were cold-anesthetized and scored for dye presence. No instances of dye transfer between painted and unpainted females were recorded (Supplemental Data Table 3).

**Figure 1.**
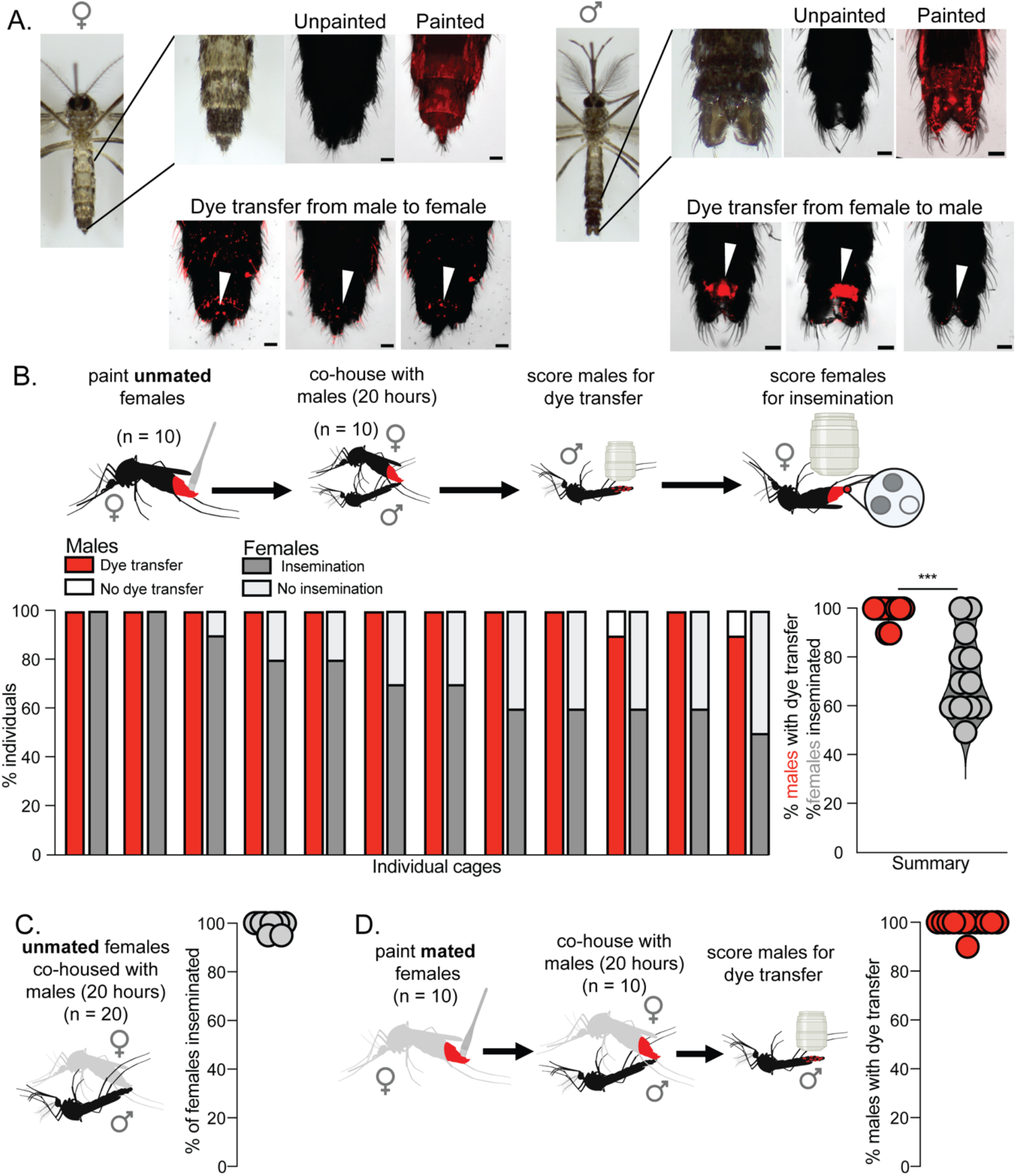
Dye transfer from females to males. **(A)** Images of a female (left) and male (right) mosquito. Top rows from left to right: a photograph of an unpainted mosquito, a photograph of the most posterior abdominal segments, and images of unpainted and painted mosquitoes. Bottom row: examples of dye transfer to unpainted females (left) and males (right) from painted partners (scale bar = 100 μm). White arrowheads indicate specific areas of transfer to the genital region. **(B)** Dye transfer and insemination between painted, unmated females and unpainted males. Dye was applied to unmated females who were co-housed with unpainted males for 20 hours. Males were then scored to determine rate of dye transfer (98.3 ± 3.89%) and female spermathecae were scored to determine rates of insemination (73.3 ± 16.7%). (n = 12 replicate cages; 10 males and 10 females/replicate). *** = p < 0.01 Mann Whitney test (U = 15, p = 0.0002). **(C)** Insemination rates during 20 hours of co-housing (99.0 ± 2.1%, n = 20 females). Unmated females were co-housed with males for 20 hours. Female spermathecae were dissected and scored for insemination (10 replicate cages; 20 females/replicate). **(D)** Dye transfer rates between painted, mated females and unpainted males (99.2 ± 2.8%). Dye was applied to mated females who were co-housed with unpainted males for 20 hours. Males were then scored for dye transfer (n = 11 replicate cages; 10 males/replicate). Females shown in gray to indicate that they were co-housed with males prior to the assay.

To score for dye transfer in all assays, unpainted animals were cold anesthetized at 4°C and dye transfer was scored using a Leica MZ10 F Fluorescence Stereomicroscope. Animals were scored as positive if fluorescent signal was detected on the genitals or abdomen. Representative images for dye transfer were taken on a Nikon Ti2-E Inverted Fluorescent Stereomicroscope in the Cy3 channel (Figure 1A). The excitation and emission spectrum of this fluorescent dye is wide, so the Cy3channel was selected for scoring dye transfer because autofluorescence of the cuticle is minimal while signal from dye is strong in that channel, however fluorescence from the dye can also be visualized in the CFP and GFP channels.

Insemination status of females was determined by spermathecal dissection. A pair of fine-tipped forceps (Dumont #5 Biology/Inox Forceps, Fine Science Tools 11252-20) were used to separate the last two abdominal segments from the rest of the abdomen, exposing the spermathecae. The spermathecae were then examined for the presence of sperm, and females were scored as inseminated if sperm was detected in the spermathecae. In all inseminated females, 2 of 3 spermathecae contained sperm, always the larger medial spermatheca and one smaller lateral spermatheca, in line with previously established patterns of sperm distribution in the spermathecae of mated *Ae. aegypti* (Roth 1948).

### Dye-transfer behavioral assays

All behavioral assays were performed in environmental rooms at 70 - 80% relative humidity, 25 – 28°C with a photoperiod of 14 hours light: 10 hours dark. Dye-transfer assays began in mid to late afternoon for a duration of 20 hours, ending the following morning, except where otherwise noted in remating encounter experiments. Animals were housed in custom cages (216mm diameter, 181mm height) and provided 10% sucrose *ad libitum* during the assay to reduce mortality. Replicates with less than 70% survival in either sex were excluded.

### Single encounter – Transfer from females to males

To score dye transfer from females to males, dye was applied to 10 females as described above. After recovery, 10 painted females were co-housed with 10 unpainted males for 20 hours, after which males were scored for dye transfer and females were dissected and scored for insemination (Figure 1B and D). We performed 13 replicate cages with painted, unmated females, 1 of which was excluded due to mortality greater than 70% among the painted females. We performed 13 replicate cages with painted, mated females, none of which were excluded.

### Single Encounter – Transfer from males to females

To score dye transfer from males to females, dye was applied to 1 male as described above. After recovery, the single painted male was co-housed with 10 unpainted females for 20 hours, after which females were scored for dye transfer and then dissected and scored for insemination (Figure 2). We performed 14 replicate cages with unpainted, unmated females, 4 of which were excluded due to death of the single painted male. We performed 14 replicate cages with unpainted, mated females, 4 of which were excluded due to death of the single painted male. Female mortality was below our threshold for exclusion in all painted male replicates.

**Figure 2.**
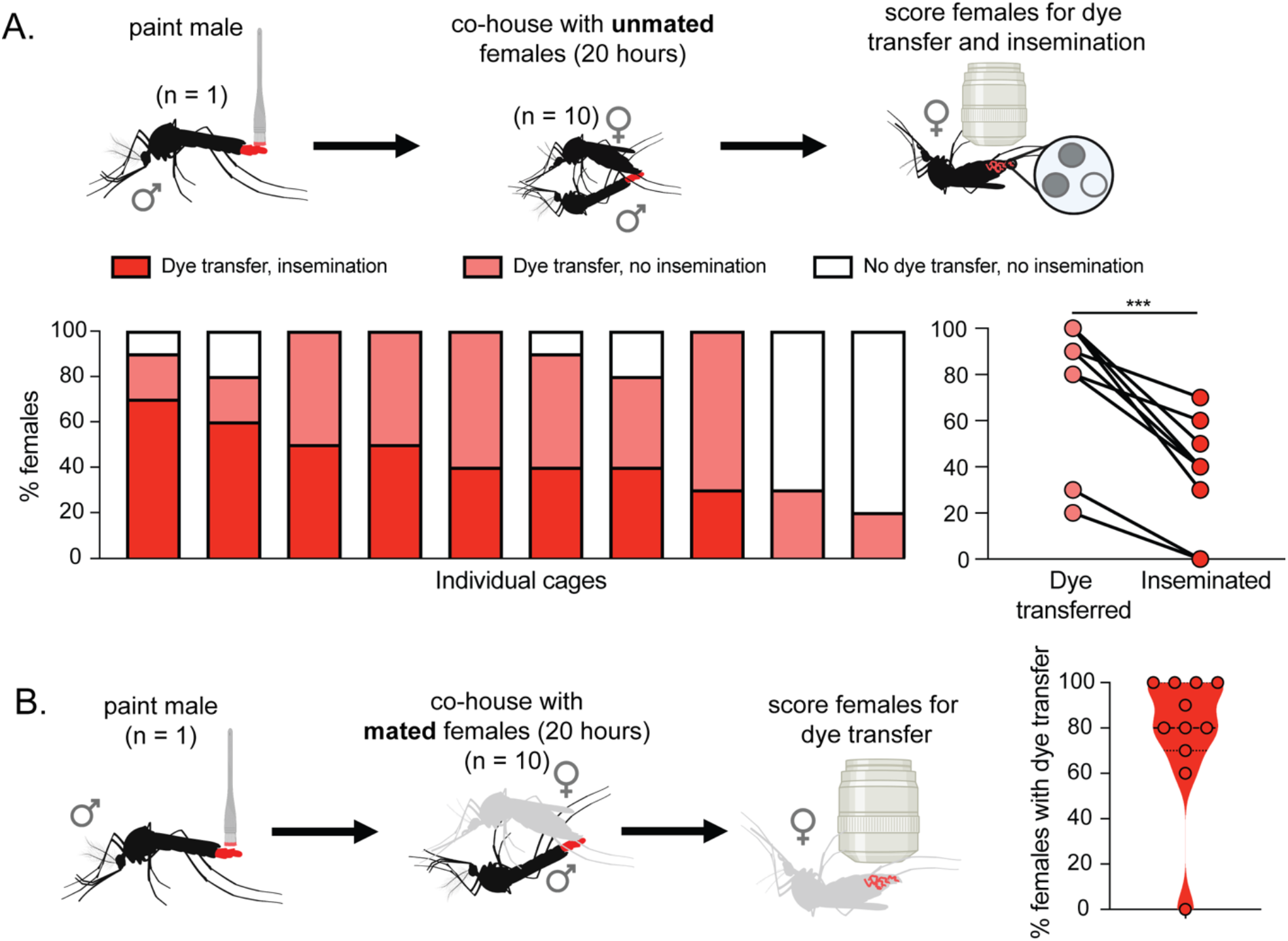
Dye transfer from male to females. **(A)** Dye transfer and insemination between a single painted male and unpainted, unmated females. An individual painted male was co-housed with unpainted, unmated females for 20 hours. Females were then scored to determine rate of dye transfer (79 ± 29.6%) and insemination (38.0 ± 21.8%) (10 replicate cages; 10 females/replicate). *** = p < 0.01, Mann Whitney test (U = 15.5, p= 0.0075). **(B)** Dye transfer between a single painted male and unpainted, mated females. An individual painted male was co-housed with unpainted, mated females for 20 hours. Females were then scored to determine rates of dye transfer (86.0 ± 12.3%) (n = 11 replicate cages; 10 females/replicate).

### Remating encounter

To investigate how attempted mating interactions may differ between initial mating encounters and subsequent encounters, unmated females were given the opportunity to mate with successive males of different genotypes. To evaluate dye transfer during instances of remating, we utilized females mutant for *npylr1*, which have been previously shown to mate with multiple males if they are encountered within hours (Duvall et al. 2017). In these assays painted, unmated females were sequentially exposed to 2 groups of unpainted males whose offspring can be genetically distinguished. Group 1 males were co-housed with females for 90 minutes; this duration was chosen because it falls within the window of HP-I/NPYLR1 paternity enforcement. Group 1 males were removed from the cage and replaced with group 2 males, which were co-housed with females for 20 hours. Males were scored for dye transfer after each encounter and offspring paternity was assigned for each female by individually genotyping offspring (Figure 3A).

**Figure 3.**
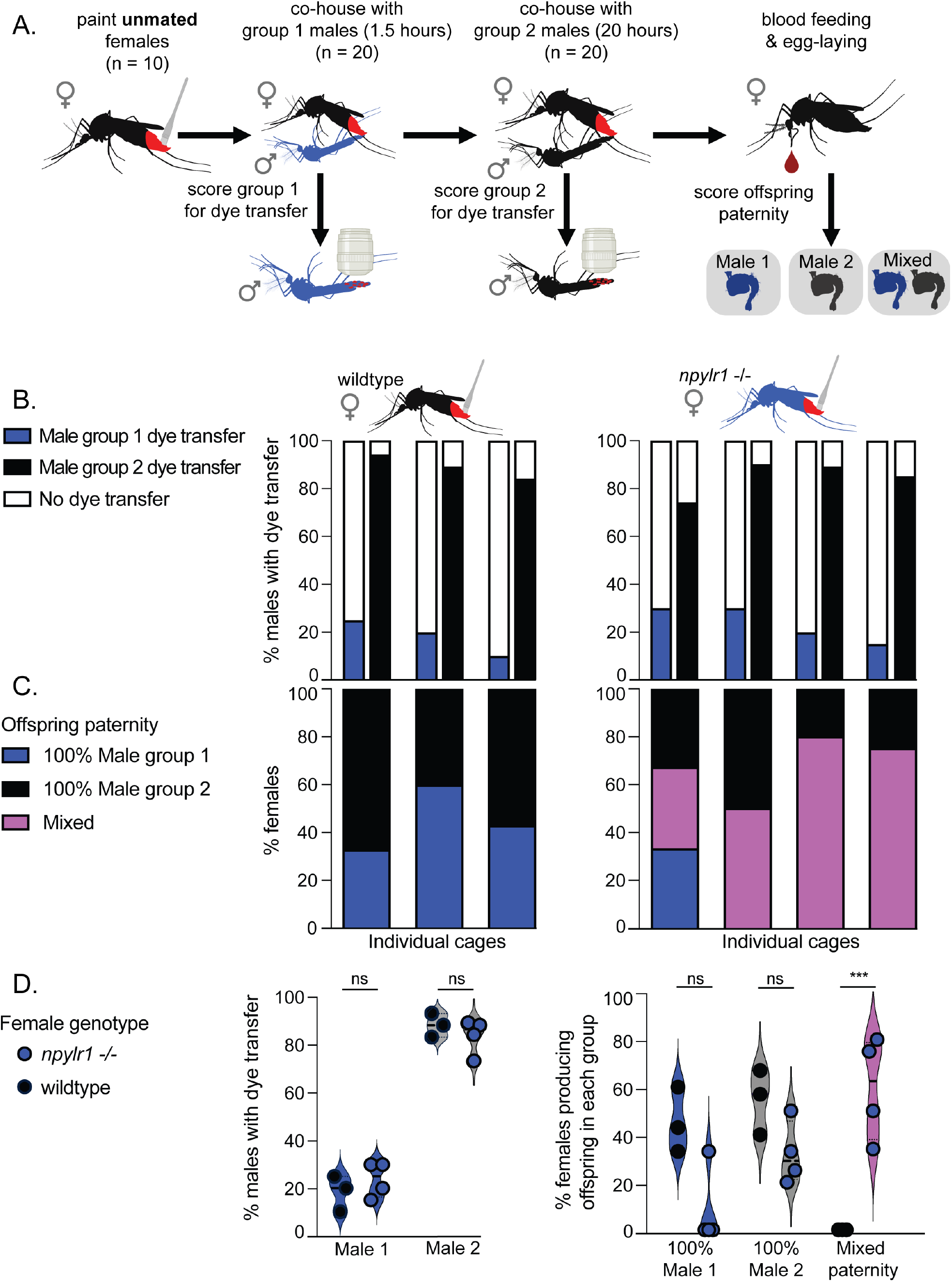
Dye transfer and paternity outcomes during remating. **(A)** Dye was applied to unmated female mosquitoes, which were co-housed with group 1 males (*npylr1-/-*, blue) for 90 minutes. Group 1 males were removed and replaced with group 2 males (Orlando, black) and co-housed for 20 hours. Males were scored for dye transfer immediately after removal. Offspring were collected individually and each female is categorized as bearing group 1 male’s offspring, group 2 male’s offspring, or mixed paternity (some offspring from each) based on genotypic differentiation of the *npylr1* locus. (**B)** Rates of dye transfer from wild-type (Orlando) females (18.3 ± 7.6%, 3 replicate cages; 20 of each male/replicate) and *npylr1-/-* females (23.8 ± 7.5%, n = 4 replicate cages; 20 males/replicate) to group 1 male (*npylr1-/-*) (84.5 ± 7.3%), and group 2 male (89.0 ± 5.0%) (Orlando). **(C)** Percent of wild-type (Orlando) females (n = 3 replicate cages; 5 – 7 females/replicate; n = 10 – 55 offspring/female) bearing offspring exclusively from group 1 male (45.3 ± 13.7%, n = 3) and group 2 male (54.7 ± 13.7%, n = 3) and percent of *npylr1-/-* females (n = 4 replicate cages, 5 – 8 females/replicate; n = 11 – 71 offspring/female) bearing offspring from group 1 male (8.3 ± 16.5%, n = 4), group 2 male (32.0 ± 13.1%, n = 4) or mixed paternity offspring (59.8 ± 21.6%, n = 4). No wild-type (Orlando) females bore offspring of mixed paternity. Bars in **(B)** and **(C)** are stacked vertically, representing dye transfer to males and paternity outcomes for the same cage of animals. **(D)** shows summary data. *** = p < 0.001, Two way ANOVAs analyzing the impact of genotype on both rates of dye transfer to group 1 males and group 2 males (F (1, 10) = 0.01451, P = 0.9065), and rate of mixed paternity offspring (F(2,15) = 20.16, P < 0.0001).

To establish a baseline mating rate of painted, unmated females exposed to males for 90 minutes, dye was applied to 10 unmated wild-type (Orlando) or *npylr1*-/-females (Liesch et al., 2013) as described above. After recovery, 20 males were introduced to the females’ cage. After 90 minutes, these males were removed and scored for dye transfer. The females were cold-anesthetized at 4°C for 10 minutes, then placed on ice and scored for insemination.

To score dye transfer from females to successive males, dye was applied to 10 unmated females as described above. After recovery, 20 unpainted males (group 1) were introduced to the females’ cage for 90 minutes. At the end of 90 minutes, all males were removed, and 20 new unpainted males (group 2) were introduced into the cage for 20 hours. In these assays, group 1 males were *npylr1-/-*, which do not show mating deficits and were used because their offspring can be unambiguously differentiated from wild-type (Orlando), which were used as group 2 males. A sex ratio of 2 males to 1 female was chosen to account for the reduced time that the females had with group 1 males (90 minutes) compared with other assays (20 hours).

We performed 4 replicates with painted, unmated *npylr1-/-* females, none of which were excluded, and 3 replicates with painted, unmated wild type (Orlando) females, none of which were excluded.

### Offspring paternity assessment

For paternity assessment, females were fed a blood meal of defibrinated sheep blood (Hardy Diagnostics DSB100) supplemented with adenosine 5’-triphosphate (ATP, 200mM in aqueous NaHCO_3_), which serves as a phagostimulant. The meal was delivered using a Hemotek artificial membrane feeder (Hemotek Ltd. SP6W1-3), and females were allowed to feed to repletion. After feeding, each replete female was individually housed in a wide fly vial with 5mL of deionized water and a 55mm-diameter Whatman filter paper cone egg-laying substrate, as in Duvall et al., 2017. Females were allowed 7 days after feeding to oviposit. After ovipositing, each female was removed, and the egg paper was pulled out of water to prevent premature hatching. After 7 days, each egg paper was placed in a separate plastic cup and hatched in 10mL of hatch broth (see Rearing above). Offspring were collected for genotyping at the pupal stage, or L4 in the case of a small number of animals that were developing slowly to genotype them alongside their siblings.

Females that did not feed to repletion or that produced fewer than 10 viable offspring were excluded from analysis. In the first replicate with wild-type females, 1 unfed female and 3 females that laid fewer than 10 eggs were excluded. In the second replicate with wild-type females, 1 unfed female and 2 females that laide fewer than 10 eggs were excluded. In the third replicate with wild-type females, 2 unfed females and 3 females that laid fewer than 10 eggs were excluded. In the first replicate with *npylr1-/-* females, 1 unfed female and 3 females that laid fewer than 10 eggs were excluded. In the second replicate with *npylr1-/-* females, 1 female that laid fewer than 10 eggs was excluded. In the third replicate with *npylr1-/-* females, 1 unfed female and 1 female that laid fewer than 10 eggs were excluded. In the fourth replicate with *npylr1-/-* females, 4 females that laid fewer than 10 eggs were excluded.

The offspring of each female were then individually genotyped using PCR and gel electrophoresis to determine their paternity. DNA extraction was performed from whole L4 larvae or pupae using Phire Tissue Direct (ThermoFisher Scientific F170), and paternity was determined by PCR-amplification of the *npylr1* locus that differentiates group 1 males (*npylr1-/-*) from group 2 males. Presence of the wild-type *npylr1* allele was detected with *npylr1* forward and reverse primers (*npylr1* forward primer: 5’-TAATCGTGTGGACTAGAAGAGGG-3’, *npylr1* reverse primer: 5’-AGCTCTTCGCAGTAGAATGTACG-3’). Presence of the mutant *npylr1-/-* allele was detected with the *npylr1* reverse primer and a forward primer embedded in the large polyubiquitin insert in the mutant (polyubiquitin forward primer: 5’-CGACTAACAGACACAAGCAC-3’; reverse primer as above). PCR products of the expected size in the agarose gel were Sanger sequenced (Genewiz) to confirm the presence of the wild-type or mutant allele.

### Analysis

GraphPad Prism was used to perform statistical analysis to assess the significance of observed results and generate associated graphs. The alpha level for all statistical analysis was set at 0.05. A Mann Whitney test was utilized to compare overall rates of dye transfer and insemination in single encounter assays involving painted males, as well as rates of insemination in wild-type (Orlando) and *npylr1*-/-females in 90 minute cohousing control assays. Two-way ANOVA analyses were conducted to examine the effects of female genotype on both the rates of dye transfer from different groups of males in remating experiments, and the proportion of females with mixed paternity offspring.

## RESULTS

### Dye transfer from females to males

Males were able to mate successfully with receptive females when dye was applied to the females, and nearly all males attempted to mate with females regardless of receptivity. In single encounter assays involving painted, unmated females and unpainted males (n = 12 replicate cages), the rate of insemination was 73.3 ± 16.7%, while 98.3 ± 3.9% of males had dye transfer from females (Figure 1B), indicating that males will attempt to mate with, and can successfully inseminate females regardless of dye application. Mann Whitney analysis showed that the proportion of males with dye transfer was significantly higher than the proportion of females that were inseminated (U=15 p=0.0002). In control experiments, we confirmed that neither incidental transfer from painted mosquitoes to the walls of the cage and then to unpainted mosquitoes, nor female to female transfer was observed (Supplemental Data Table 1 and 3).

Female *Ae. aegypti* are non-receptive to males after they have successfully mated (Gwadz et al. 1971). We find that rates of insemination in control assays are maximal after 20 hours of cohousing with males; 99.0 ± 2.0% of females are inseminated at this timepoint in control experiments (Figure 1C), confirming that 5 days of co-housing provides ample time to ensure that females are mated before the assay. In single encounter assays with painted, previously-mated females and unpainted males (n = 12 replicate cages), mated females transfer dye to 99.2 ± 2.8% of males, despite mated females being unreceptive and refractory to subsequent mating (Gwadz et al. 1971) (Figure 1D).

### Dye transfer from males to females

Painted males were able to mate with and inseminate receptive females, and males attempted to mate with females regardless of receptivity. In single encounter assays involving painted males and unmated, unpainted females (n = 10 replicate cages), 79.0 ± 29.6% of females had dye transfer and 38.0 ± 21.8% of females were inseminated (Figure 2A). In each cage, all inseminated females had dye transfer and in each cage dye transfer exceeded insemination. A Mann Whitney test revealed that the overall level of dye transfer to females was significantly higher than insemination (38.0 ± 21.8%) (U = 15.5, p= 0.0075.) (Figure 2A), indicating that a single male will make contact with more females than he will successfully inseminate.

Single-encounter assays with a single painted male housed with unpainted, previously-mated females (n = 11 replicate cages) resulted in dye transfer to 86.0 ± 12.3% of females. In one case, the male did not transfer dye to any females (Figure 2B), indicating either that the male did not attempt to mate or that the females rejected this male before he was able to make contact.

These data show that males frequently make dye-transferring contact with both unmated and previously mated, unreceptive females. This indicates a male will attempt to mate, regardless of the female’s receptivity, consistent with previous studies (Gwadz et al. 1971; Jones and Pilitt 1973).

### Dye transfer during remating

In order to determine whether rates of dye transfer differ between wild-type (Orlando) females and those with disrupted mating pathways we scored interactions between wild-type (Orlando) and *npylr1* mutant females who sequentially encountered two groups of males. Rates of dye transfer to group 1 males during the 90 minute exposure window were 18.3 ± 7.6% (n = 3 replicate cages) in wild-type (Orlando) and 23.8 ± 7.5% (n = 4 replicate cages) in *npylr1-/-* females. Rates of dye transfer to group 2 males were 89.0 ± 5.0% (n = 3 replicate cages) from wild-type females and 84.5 ± 7.3% (n = 4 replicate cages) from *npylr1-/-* females (Figure 3B). A two-way ANOVA showed that in both wild-type (Orlando) and *npylr1-/-* females, the rates of dye transfer to group 1 males were significantly lower than rates to group 2 males (F (1, 10) = 298.2, P<0.0001) but that female genotype did not impact rates of transfer (F (1, 10) = 0.01451, P=0.9065) (Figure 3D). Mann Whitney analysis showed that within a 90-minute window, insemination rates of painted *npylr1* mutants (21.3 ± 20.8%, n = 3 replicate cages) and wild-type (*Orlando*) females (32.5 ± 19.1%) were not significantly different (U=3.0, p=0.7) (Supplemental Data Table 2). These data indicate that wild-type (Orlando) and *npylr1-/-* females make comparable dye-transferring contacts with both groups of males.

To score offspring genotype, females were blood fed and housed in individual vials for oviposition so that offspring could be attributed to a single female. Wild-type (Orlando) females produced offspring fathered exclusively by group 1 (45.3 ± 13.7%) <u>or</u> group 2 (54.7 ± 13.7%) but never produced any mixed-paternity offspring, despite dye-transferring interactions with both males (n = 3 replicate cages) (Figure 3C and 3D). Although some *npylr1* mutant females produced offspring fathered exclusively by group 1 (8.3 ± 16.5%) or group 2 (32.0 ± 13.1%), this group also included individual females who bore offspring of mixed paternity within a single clutch (59.8 ± 21.6%) (n = 4 replicate cages), a phenomenon that was never observed in the wild-type females (Figure 3C and 3D). Two-way ANOVA analysis showed that *npylr1* mutants have significantly more mixed paternity offspring than wild-type (Orlando) females (F(2,15) = 20.16, P < 0.0001). These data indicate that although both wild-type (Orlando) and *npylr1* mutant females make similar copulatory contacts both groups of males, <u>only</u> *npylr1* mutant females are ever successfully inseminated by and produce offspring fathered by both group 1 and group 2 males.

## DISCUSSION

In this study, we utilize a hydrophobic oil-based fluorescent dye to score physical interactions between males and females to assess attempted and successful mating in *Ae. aegypti*. We show that both painted males and females transfer dye to the posterior abdominal segments and areas immediately surrounding the genitals of unpainted animals of the opposite sex, whether the female is receptive or not. The absence of insemination without dye transfer validates the method as faithfully reporting attempted mating. Previous observations have shown that a single male mosquito can inseminate 5 to 7 females before exhausting his sperm, which is in line with our observations (Gwadz et al. 1971; Jones 1973). Our observation that a single painted male always transfers dye to more receptive females than he will successfully inseminate also suggests that male mosquitoes continue to attempt to mate when given the opportunity, even if they have depleted their sperm stores.

Although female rejection has been characterized as flying away from the male, abdominal twisting, flicking of wings, kicking males and/or failure to open vaginal plates and extrude ovipositor, we lack a mechanistic understanding of how and when these behaviors are deployed (Roth 1948; Gwadz et al. 1971; Jones and Pilitt 1973; Cator and Harrington 2011). The presence of dye transfer to the genitalia of the opposite sex, especially in the case of unreceptive females, supports the observation of “pseudocopulation” reported in Gwadz 1971, where there was genital contact, but no transfer of semen. These observed high rates of attempted mating culminating in contact of the external genitalia suggests the importance of mechanisms of rejection once animals reach the point of genital contact and imply additional methods of mate rejection on the part of unreceptive females. Such methods include external barriers such as closing of the vaginal sclerite to prevent insemination, or sperm rejection after copulation before the sperm reach the spermathecae.

There are limitations to this method, which must be considered when designing assays. Our assays were not designed to identify pair-wise mating interactions, although individual housing or combinatorial use of multiple dyes could be used to achieve this. Although dye transfer reports mating attempts in which genital contact was achieved it does not report all mating attempts; previous work indicates that many female rejection behaviors preclude physical contact and would not result in dye transfer (Aldersley and Cator 2019).

Although painted males can inseminate females and painted females can be inseminated, rates of mating in cages with painted animals were lower than in assays in which both partners were unpainted, regardless of the genotype (compare Figure 1B and 1C). It is possible that painted animals require a longer to reach maximal mating rates. This may be because dye application or cold stress leads to reluctance to mate, reduces painted females’ attractiveness to potential mates, or reduces animals’ ability to maneuver.

There was no discernable difference in spatial patterns of dye transfer between wild-type and mutant animals or receptive and non-receptive females. Dye transfer indicates attempts at mating and must be considered along with female mating status at the beginning of the assay and insemination status after the assay to determine whether mating attempts were successful.

Although we developed this assay for use in *Ae. aegypti*, it is likely to succeed across many insect species. It may offer further insight and greater resolution to researchers interested in assessing males’ attempted interspecific mating, as in the case of *Ae. aegypti* and *Ae. albopictus. Ae. albopictus* males are known to attempt to mate with females of closely-related species, including *Ae. aegypti* (Leahy and Craig 1967; Tripet et al. 2011; Bargielowski et al. 2015; Lounibos et al. 2016). Disrupting mating systems to exogenously suppress receptivity and effectively sterilize female mosquitoes is also of potential interest to mosquito control programs. A well-established method of controlling invasive and pest insect populations in the wild involves the mass introduction of sterilized or genetically modified males. The success of this approach relies on the fitness of the released males and their ability to successfully compete with wild males to mate with wild females (Alphey et al. 2010; Benelli 2015; Lees et al. 2015). The effectiveness and sustainability of these control strategies in the long run will be improved by releasing males with a high probability of mating success, which will facilitate the success of the program. Researchers could test lab strains’ ability to compete with wild populations to interact with females. It is our hope that the accessibility of our method will be of use to researchers working in a variety of insect mating systems, from cryptic mating behaviors to biological control.

## Acknowledgments

We thank members of the Duvall Lab for their critical comments on the manuscript. We thank Lauren Subramaniam & Candace Cochran for their assistance in completing assays and data collection.

## References

Aldersley A, Cator LJ. 2019. Female resistance and harmonic convergence influence male mating success in Aedes aegypti. Scientific Reports 9:2145.

Alphey L, Benedict M, Bellini R, Clark GG, Dame DA, Service MW, Service MW, Dobson SL. 2010. Sterile-insect methods for control of mosquito-borne diseases: an analysis. Vector-Borne and Zoonotic Diseases 10.

Attardo GM, Hansen IA, Raikhel AS. 2005. Nutritional regulation of vitellogenesis in mosquitoes: Implications for anautogeny. Insect Biochemistry and Molecular Biology 35:661–75.

Bargielowski IE, Lounibos LP, Shin D, Smartt CT, Carrasquilla MC, Henry A, Navarro JC, Paupy C, Dennett JA. 2015. Widespread evidence for interspecific mating between Aedes aegypti and Aedes albopictus (Diptera: Culicidae) in nature. Infection, Genetics and Evolution 36:456–61.

Benelli G. 2015. Research in mosquito control: current challenges for a brighter future. Parasitology Research 114:2801–5.

Bhatt S, Gething PW, Brady OJ, Messina JP, Farlow AW, Moyes CL, Drake JM, Brownstein JS, Hoen AG, Sankoh O, Myers MF, George DB, Jaenisch T, Wint GRW, Simmons CP, Scott TW, Farrar JJ, Hay SI. 2013. The global distribution and burden of dengue. Nature 496:504–7.

Carrasquilla MC, Lounibos LP. 2015. Satyrization without evidence of successful insemination from interspecific mating between invasive mosquitoes. Biology Letters 11:20150527.

Carvalho DO, Chuffi S, Ioshino RS, Marques ICS, Fini R, Costa MK, Araújo HRC, Costa-da-Silva AL, Kojin BB, Capurro ML. 2018. Mosquito pornoscopy: Observation and interruption of Aedes aegypti copulation to determine female polyandric event and mixed progeny. PLoS ONE 13:e0193164.

Cator LJ, Harrington LC. 2011. The harmonic convergence of fathers predicts the mating success of sons in Aedes aegypti. Animal Behaviour 82:627–33.

Clifton ME, Correa S, Rivera-Perez C, Nouzova M, Noriega FG. 2014. Male Aedes aegypti mosquitoes use JH III transferred during copulation to influence previtellogenic ovary physiology and affect the reproductive output of female mosquitoes. Journal of Insect Physiology 64:40–47.

Craig GB. 1967. Mosquitoes: Female Monogamy Induced by Male Accessory Gland Substance. Science 156:1499–1501.

Dimond JB, Lea AO, Hahnert WF, DeLong DM. 1956. The Amino Acids Required for Egg Production in Aedes aegypti. The Canadian Entomologist 88:57–62.

Duvall LB, Basrur NS, Molina H, McMeniman CJ, Vosshall LB. 2017. A Peptide Signaling System that Rapidly Enforces Paternity in the Aedes aegypti Mosquito. Current Biology 27:3734–3742.e5.

Eberhard WG. 1991. Copulatory Courtship and Cryptic Female Choice in Insects. Biological Reviews 66:1–31.

Fuchs MS, Craig GB, Hiss EA. 1968. The Biochemical Basis of Female Monogamy in Mosquitoes I. Extraction of the Active Principle From Aedes aegypti. Vol 7.

Guerbois M, Fernandez-Salas I, Azar SR, Danis-Lozano R, Alpuche-Aranda CM, Leal G, Garcia-Malo IR, Diaz-Gonzalez EE, Casas-Martinez M, Rossi SL, Del Río-Galván SL, Sanchez-Casas RM, Roundy CM, Wood TG, Widen SG, Vasilakis N, Weaver SC. 2016. Outbreak of Zika Virus Infection, Chiapas State, Mexico, 2015, and First Confirmed Transmission by Aedes aegypti Mosquitoes in the Americas. Journal of Infectious Diseases 214:1349–56.

Gwadz RW, Craig GB, Hickey WA. 1971. Female Sexual Behavior as the Mechanism Rendering Aedes aegypti Refractory to Insemination. The Biological Bulletin 140:201–14.

Helinski MEH, Valerio L, Facchinelli L, Scott TW, Ramsey J, Harrington LC. 2012. Evidence of Polyandry for Aedes aegypti in Semifield Enclosures. The American Journal of Tropical Medicine and Hygiene 86:635–41.

Hiss EA, Fuchs MS. 1972. The effect of matrone on oviposition in the mosquito, Aedes aegypti. Journal of Insect Physiology 18:2217–27.

Jones JC, Pilitt DR. 1973. Observations on the Sexual Behavior of Free Flying Mosquitoes. The Biological Bulletin 144:480–88.

Leahy MG, Craig GB. 1967. Barriers to Hybridization Between Aedes aegypti and Aedes albopictus (Diptera: Culicidae). Evolution 21:41–58.

Lees RS, Gilles JR, Hendrichs J, Vreysen MJ, Bourtzis K. 2015. Back to the future: the sterile insect technique against mosquito disease vectors. Current Opinion in Insect Science 10:156–62.

Liesch J, Bellani LL, Vosshall LB. 2013. Functional and Genetic Characterization of Neuropeptide Y-Like Receptors in Aedes aegypti. PLoS Neglected Tropical Diseases 7:e2486.

Lounibos LP, Bargielowski I, Carrasquilla MC, Nishimura N. 2016. Coexistence of Aedes aegypti and Aedes albopictus (Diptera: Culicidae) in Peninsular Florida Two Decades After Competitive Displacements. Journal of Medical Entomology 53:1385–90.

MacGregor, M.E. 1915. Notes on the Rearing of Stegomyia fasdata in London. Journal of Tropical Medicine and Hygiene 18:193–96.

Naccarati C, Audsley N, Keen JN, Kim J-H, Howell GJ, Kim Y-J, Isaac RE. 2012. The host-seeking inhibitory peptide, Aea-HP-1, is made in the male accessory gland and transferred to the female during copulation. Peptides 34:150–57.

Richardson JB, Jameson SB, Gloria-Soria A, Wesson DM, Powell J. 2015. Evidence of Limited Polyandry in a Natural Population of Aedes aegypti. The American Journal of Tropical Medicine and Hygiene 93:189–93.

Rogers DJ, Wilson AJ, Hay SI, Graham AJ. 2006. The Global Distribution of Yellow Fever and Dengue. In: Advances in Parasitology Elsevier. p. 181–220.

Roth LM. 1948. A Study of Mosquito Behavior. An Experimental Laboratory Study of the Sexual Behavior of Aedes aegypti (Linnaeus). American Midland Naturalist 40:265.

Spielman A, Leahy MG, Skaff V. 1967. Seminal Loss in Repeatedly Mated Female Aedes aegypti. The Biological Bulletin 132:404–12.

Tripet F, Lounibos LP, Robbins D, Moran J, Nishimura N, Blosser EM. 2011. Competitive Reduction by Satyrization? Evidence for Interspecific Mating in Nature and Asymmetric Reproductive Competition between Invasive Mosquito Vectors. The American Journal of Tropical Medicine and Hygiene 85:265–70.

Weaver SC, Costa F, Garcia-Blanco MA, Ko AI, Ribeiro GS, Saade G, Shi P-Y, Vasilakis N. 2016. Zika virus: History, emergence, biology, and prospects for control. Antiviral Research 130:69–80.

